# An information-theoretic perspective on the costs of cognition

**DOI:** 10.1101/208280

**Authors:** Zénon Alexandre, Solopchuk Oleg, Pezzulo Giovanni

## Abstract

In statistics and machine learning, model accuracy is traded off with complexity, which can be viewed as the amount of information extracted from the data. Here, we discuss how cognitive costs can be expressed in terms of similar information costs, i.e. as a function of the amount of information required to update a person’s prior knowledge (or internal model) to effectively solve a task. We then examine the theoretical consequences that ensue from this assumption. This framework naturally explains why some tasks – for example, unfamiliar or dual tasks – are costly and permits to quantify these costs using information-theoretic measures. Finally, we discuss brain implementation of this principle and show that subjective cognitive costs can originate either from local or global capacity limitations on information processing or from increased rate of metabolic alterations. These views shed light on the potential adaptive value of cost-avoidance mechanisms.

## 1. Introduction

Demanding cognitive tasks, such as mental arithmetic, are strongly aversive: we tend to avoid partaking in such tasks and they lead to unpleasant subjective feeling of mental exertion (Inzlicht et al., 2015). Various studies have revealed that we take into consideration a measure of cognitive cost or cognitive effort when deciding whether or not to engage in a task (Benoit et al., 2017; Kool et al., 2010; Manohar et al., 2015; Schmidt et al., 2012; Westbrook et al., 2013; Westbrook and Braver, 2015). Furthermore, prolonged performance of demanding tasks leads to cognitive fatigue, which is characterized by a subjective dimension – i.e. feeling of exhaustion, impression of worsened ability and decreased willingness to engage in mental activities (Hockey, 2011; van der Linden et al., 2003) – and an objective dimension, with an actual decrease of task performance (Bailey et al., 2007; Tanaka, 2015; van der Linden et al., 2003). However, it is still unclear what is the origin of cognitive costs (i.e., what is costly about cognitive processing?), how to specify them quantitatively, and whether cognitive costs and cognitive fatigue have some adaptive value.

## 2. The information cost of cognitive processes

Recent advances in artificial intelligence and computational neuroscience have led to formalization of cognition as a bounded rationality process (Friston, 2010; Kingma and Welling, 2013; Ortega and Braun, 2013; Tishby et al., 2000; Tkačik and Bialek, 2014). According to this view, rather than aiming systematically at the optimal solution to computational problems, cognitive processes trade off performance with computational costs. Remarkably, the way computational costs are formalized across these different studies is very consistent, despite their different approaches. In fact, whether one starts from an inference problem, in which the evidence for a model is maximized given some data (Genewein et al., 2015; Kingma and Welling, 2013; Tishby et al., 2000), or whether one is more generally attempting to minimize the entropy of future states (Friston, 2010), or whether one takes a decision making perspective, in which expected utility is maximized (Ortega et al., 2015), or even from the point of view of thermodynamics (Ortega and Braun, 2013; Sengupta et al., 2013), information cost is framed as a measure of divergence between an initial belief (or prior probability distribution over a variable of interest *x*, such as expected reward) and an updated belief (or posterior probability distribution over the same variable *x*) obtained after receiving new data. This measure of difference between probability distributions, called the Kullback-Leibler (KL) divergence, represents the amount of information one needs to collect in order to update the prior to the posterior (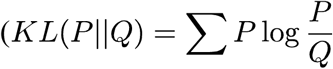 for probability distributions *P* and *Q*). Here, “amount of information” is meant in the sense of Shannon’s definition of information in terms of *surprisal* or the negative log probability of the data (Shannon, 1948). In other words, expected data provides little information while unexpected data is very informative. If one wants to encode such data without error, the number of binary symbols that will be needed is at least equal to the average surprisal, or *entropy* of the data (entropy is often represented by the letter *H*: *H*(*P*) − Σ *P* log _2_ *P*).

Whether we are considering an inference problem (identifying latent causes of observations) or a decision making problem (deciding what to do), cognitive activity can be viewed as a situation in which input data allows us to refine our previous assumption (about the most probable cause or the best action) to a new, more accurate belief and the cost of this process corresponds to the reduction in the entropy that it causes.

The question of the cost of cognitive control – i.e. the selection of appropriate behaviour in the face of environmental stimulation, on the basis of internal goals (Miller and Cohen, 2001; Pezzulo et al., 2018) - has been addressed for more than 50 years, pioneered, separately, by Hick and Hyman (Hick, 1952; Hyman, 1953). In these early works, cognitive cost was framed in terms of the entropy of the response choice (we will refer henceforth to response choices by the variable name *y*): *H(p(y))*. A typical example is the digit-key association task, in which participants have to press the key that corresponds to the digit they see on the screen. In this task, participants need to update their prior distribution of responses *p*_*0*_*(y)* to a posterior distribution *p(y*|*x)* (read as the probability of *y* given *x*, where *x* refers to the input data) in which all the probability mass is in one stimulus-response association (i.e., one cell becomes 1 and all the other cells become zero, see Figure 1, dark grey). If there are 4 possible digits and their probabilities of occurrence are equal, the KL divergence between *p*_*0*_*(y)* and *p(y*|*x)* is equal to *H(p*_*0*_*(y))*, which in the present case is log(4). Strikingly, what Hick and Hyman showed is that reaction time is a linear function of *H(p*_*0*_*(y)) (Hick, 1952; Hyman, 1953)*, confirming that it is an accurate measure of cognitive cost in this context and suggesting that the rate of information for this task is a constant (i.e. the number of bits processed per unit of time is constant).

**Figure 1.**
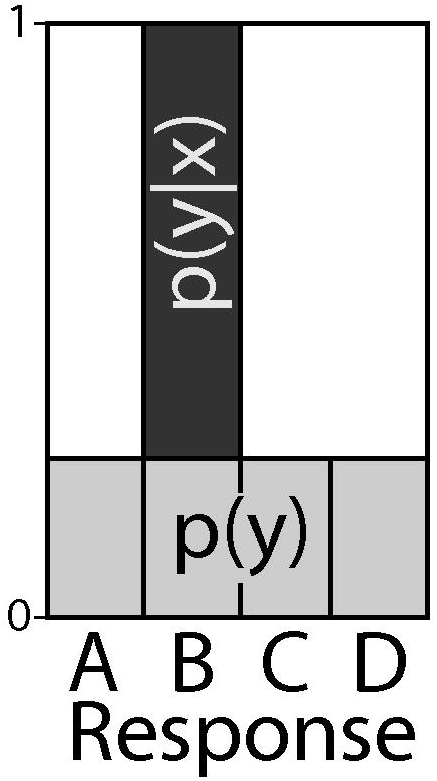
Probability distribution of responses in a simple digit-key association task, before (light gray) and after (dark gray) seeing the to-be-pressed digit on the screen.

However, the uncertainty in the response is clearly not the sole determinant of task complexity. In the task above, if one varies the number of stimuli associated to each button press while keeping the number of responses constant (e.g. 2 stimuli become associated to the same button presses, leading to 8 stimuli for 4 button presses), *H(p*_*0*_*(y))* remains constant but reaction times increase, and they do so linearly with *H(p*_*0*_*(x))* (Wifall et al., 2016). A similar increase in reaction time was reported in other tasks in which the complexity of the stimulus varies, while the number of response choices remains constant (Fan, 2014; Fan et al., 2008). To explain these findings, it is necessary to extend the aforementioned information theoretic framework, such that cognitive costs depend on both *p(x)* and *p(y)*.

It is worth noting that *x* stands for sensory data coming from the outside world and not an internal variable of the agent – hence it cannot be used directly for our formalization. For this, we need to introduce a novel, auxiliary variable *z* that would stand for the internal representation of *x*, which the system uses to choose the action *y*: *x* → *z* → *y*. The addition of this intermediate variable *z* affords different levels of compression of the input *x* (*z* can extract more or less of the information that is available in *x*), which is an important feature for a model of cognition (Grau-Moya and Braun, 2015; Park and Pillow, 2017; Tishby et al., 2000).

Therefore, information cost 𝒞 becomes the KL divergence between the prior internal representation of the input *p*_*0*_*(z)* and its posterior after observing the outside world *p(z*|*x)*, to which we need to add the KL divergence between the prior distribution of responses *p*_*0*_*(y)* and their posterior distribution *p(y*|*z)*:

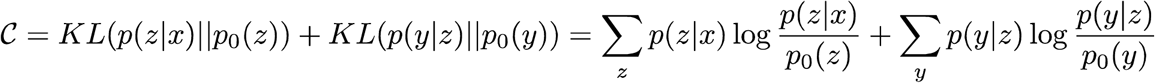

It is worth adding that if one considers the average of these information costs across many trials, and assuming usage of an optimal, marginal prior (see below and (Tishby et al., 2000)), the total information cost becomes:

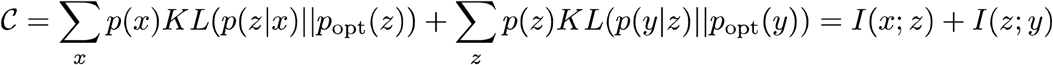

where *I(x;z)* refers to the *mutual information* between *x* and *z*, which indicates the reduction in the entropy of *z* after observing *x* (which is closely related to the notion of epistemic value in active inference (Friston et al., 2015)), while *I(z;y)* represents the reduction in entropy of the response given some internal representation of the input *z*. We can see from the above formula that this framework implements the property of 𝒞 that we wanted: its dependence on both the complexity of the input data *z* and the complexity of the responses *y*.

Finally, in order to account for the cognitive costs associated with classic tasks such as the Stroop task for example, we need to make one last addition to our framework, in agreement with earlier proposals by Koechlin and Summerfield (Koechlin and Summerfield, 2007). In the Stroop task, subjects must either read a colour word or name the colour of the ink with which the word is written. When both sources of information are incongruent (e.g. the word blue is written with red ink), it is more difficult to name the ink colour than to read the word (MacLeod, 1991). This indicates that the stimulus triggers an action (reading the word) that is independent of the task context and that interferes with the response instructed by the task (naming the ink colour) (Cohen et al., 1990). In order to account for this, we need to add a new variable *T* that represents the context of the task. The stimulus then triggers a default, automatic conditional distribution of responses *p(y*|*z)* (that is thus independent of context), which is finally updated to the final, context-dependent distribution: *p(y*|*z,T)*. The optimal automatic response distribution is the marginal of the final response distribution (i.e. the final distribution averaged over all contexts): 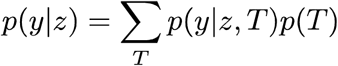.

Our final formula for total information cost with optimal priors becomes (see also Figure 2):

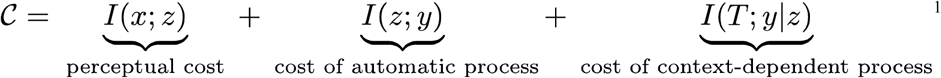

**Figure 2.**
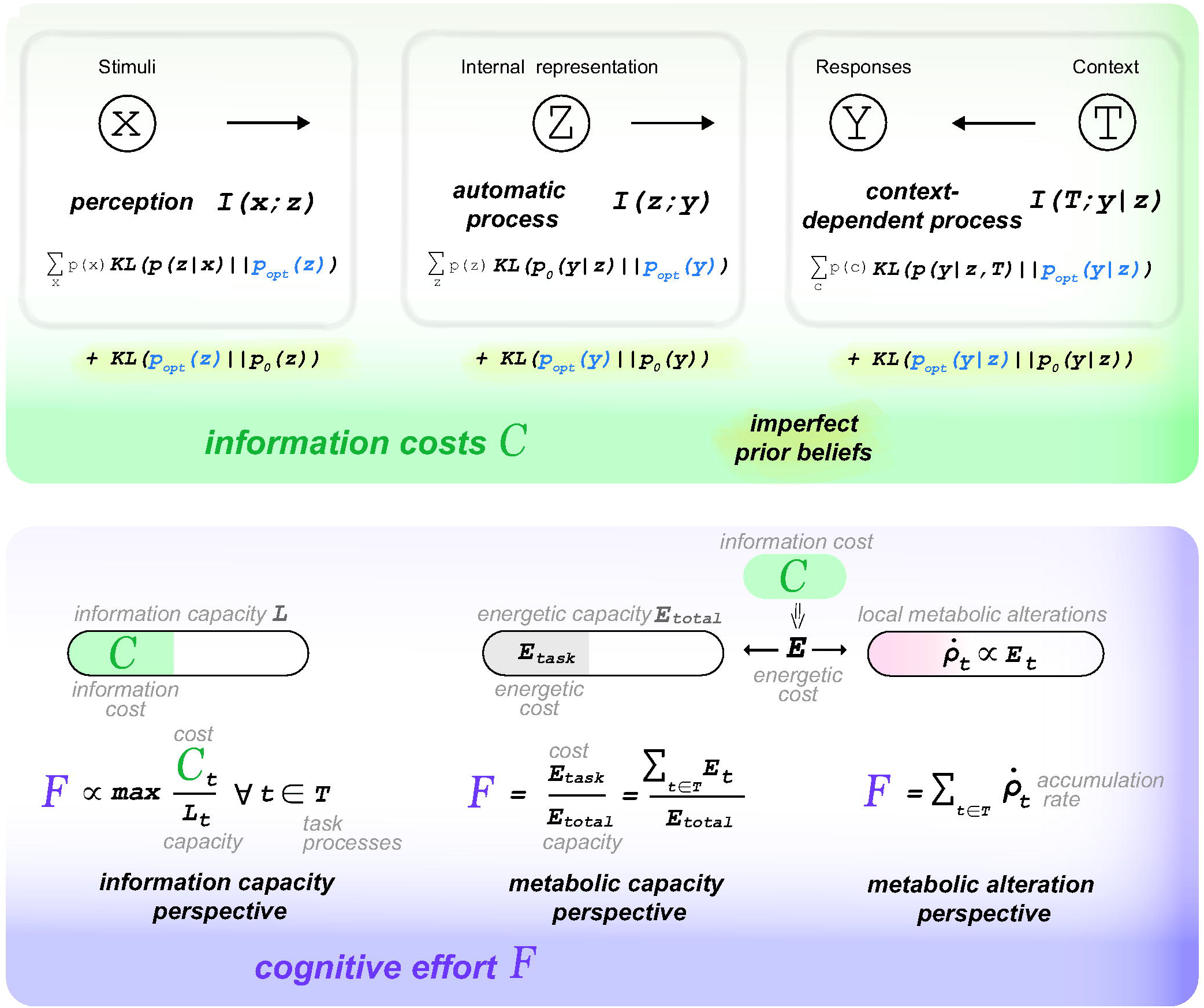
Summary of the present theoretical framework. The upper panel, shaded in green, illustrates the mathematical formulation of information costs *C*. The lower panel, shaded in blue, shows the three proposals that we make regarding how these information costs *C* could translate into subjective effort *F*.

## 3. Predictions of the framework

So far, we have shown that information cost of a cognitive task can be framed as the sum of three terms: the mutual information between inputs and their internal representations; the mutual information between internal representations and automatic responses; and the mutual information between contextual information and automatic responses. Now we explain in more details how to apply this framework in practice and discuss its predictions in terms of expected cost of different types of tasks. This theory predicts that certain kinds of tasks - those that have *many degrees of freedom*, are *unfamiliar*, necessitate to go *against natural biases*, have *variable statistical structure* or *low signal to noise ratios* - will lead to large information costs.

### 3.1 The costs of tasks that have many degrees of freedom

Tasks that have *many degrees of freedom*, or equivalently, a wide probability distribution of state-action combinations, are expected to be cognitively costly under the proposed information theoretical framework, as they imply low, widely spread prior probabilities - and thus significant information costs to update the priors (see Figure 3). Arithmetic tasks, chess games or creative writing, which are well known for being cognitively demanding (Hess and Polt, 1964; Kellogg, 1987; Marshall, 2002; Westbrook and Braver, 2015b) all assign a small prior probability mass for each possible decision and hence, lead to large divergence with the final posterior obtained when the choice has been made. Intuitively, this would be equivalent to having a very wide response space in Figure 1, with the same, small probability in each cell. Similarly, tasks that demand a deep contextualisation of the stimulus-response associations (e.g. learning to navigate in a complex maze) will lead to multidimensional prior distributions whose space can inflate very fast, leading also to fast increase in complexity.

**Figure 3.**
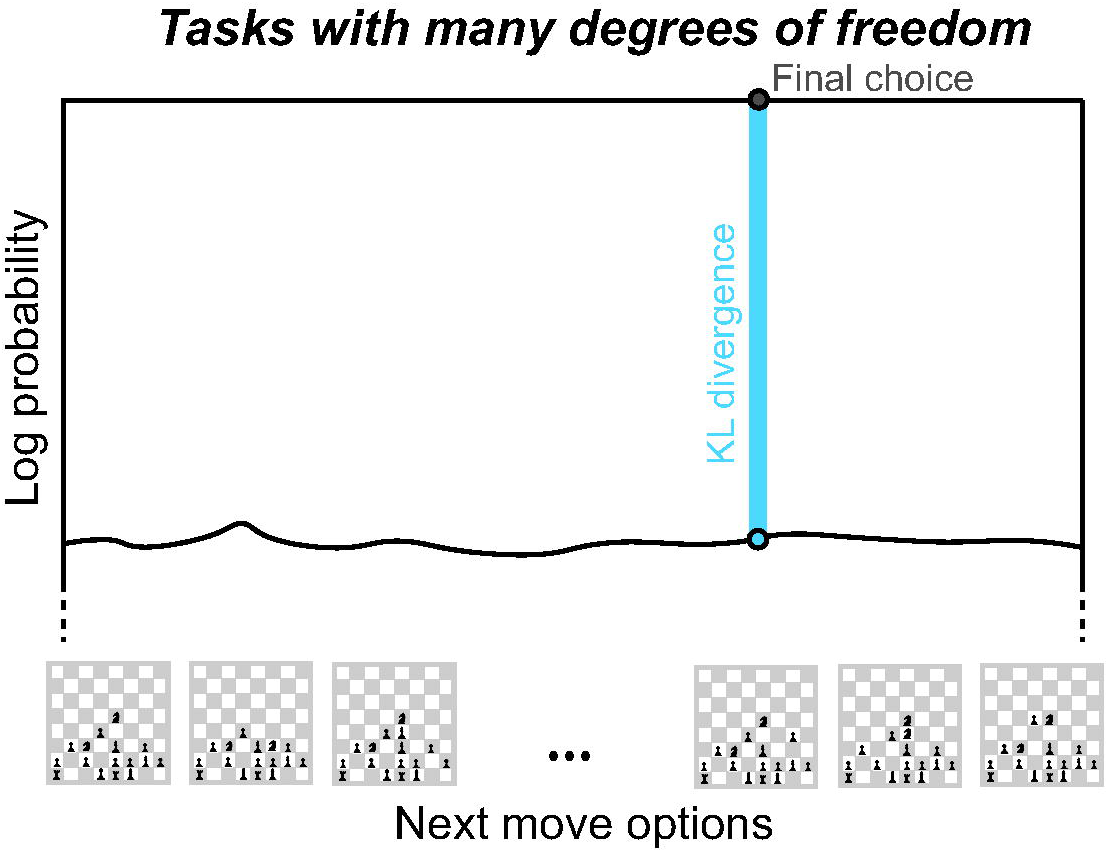
Schematic illustration of the probability distribution of different chess moves. The large size of the state-action space leads to small prior probabilities for all options. The information cost associated with the selection of the final choice is shown as a thick blue line, which represents the KL divergence between the prior and the posterior probability distribution of the chess moves. Log probabilities are shown on the y-axis for consistency with the mathematical definition of the KL divergence, such that for the chosen action, the sum of its log prior probability *log(P*_*0*_) and KL equals 0 (since KL equals to *-log(P*_*0*_), given that *p(y*|*z,T)* for the chosen action is equal to one).

Interestingly, the complexity of the environment *x* has no impact on cognitive costs. Only the complexity of *z*, its internal representation, will affect information cost 𝒞. This can be understood intuitively if one considers, for example, that during the digit-key association task, the digit presented on the screen could be represented as an image of arbitrary size and complexity. Indeed, all *z* needs to encode about *x* is the identity of the digit (i.e. 1 to 4) and what will matter for task difficulty is only the probability of occurrence of that digit (see above). An interesting implication is that – as known since the beginning of artificial intelligence – the way one encodes or represents the task drastically affects the complexity (or event the possibility) of solving it (Minsky, 1961; Simon, 1956). We will return to the issue of information compression below.

### 3.2 The costs of novel or unfamiliar tasks

A similar issue arises with *unfamiliar tasks*, that is, tasks in which the statistical structure of the sensory states, state transition probabilities (conditional on the performed actions) or action policies (e.g. sequences of motor actions) are poorly known (see Figure 4). This lack of knowledge of statistical properties of the task leads to non-optimal encoding and large information costs. This is because participants will have to start from uninformative prior distributions *p*_*0*_*(z), p*_*0*_*(y)* and *p*_*0*_*(y*|*z)* that may be far from the true marginal distributions of the task, which they will have to learn across trial repetitions (Genewein et al., 2015; Tishby et al., 2000). In this case the additional cost of starting with the wrong priors can be formalized

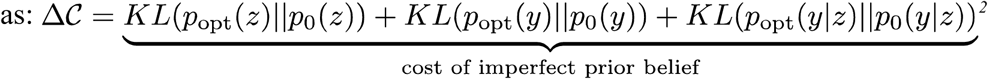

**Figure 4.**
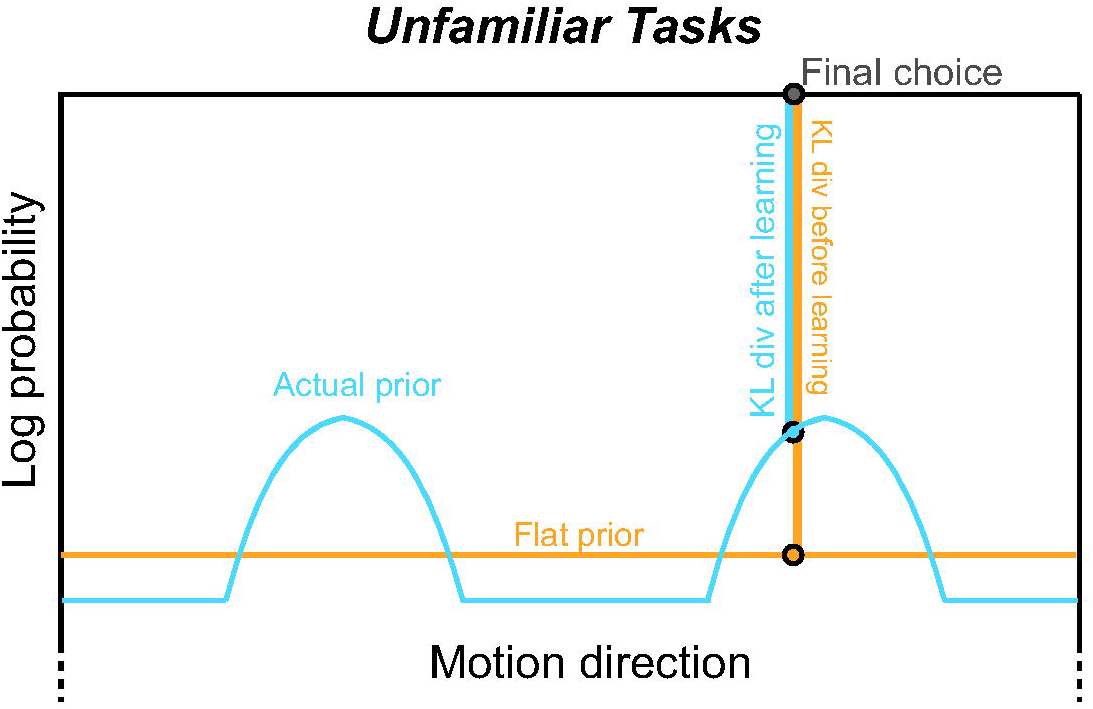
Example of the difference in information costs between learned and novel task contingencies. The example is inspired by the (Chalk et al., 2010) study, where subjects learn to perform a motion discrimination task in which motion direction is distributed non-uniformly.

Importantly, here we are assuming that participants have understood the rules of the task and know how to perform it correctly. Therefore, this additional cost of imperfect priors represents the extra information participants need to process because of their poor assumption of task statistics - *p*_*0*_*(z), p*_*0*_*(y)* and *p*_*0*_*(y*|*z)* - not the cost of learning the task rules. This corresponds to the notion of *cross-entropy* in information theory: the number of symbols needed to encode data when using the wrong encoding scheme - i.e. one that is based on the wrong probability distribution - is always larger than entropy (the number of symbols necessary when the correct distribution is assumed). In behavioural terms, this corresponds to the well-known effect of training on reaction time (Teichner and Krebs, 1974), subjective effort (Mykityshyn et al., 2002), or pupil size (Hyönä et al., 1995; Recarte and Nunes, 2000; Solopchuk et al., 2016), regarded as a reliable index of effort (Beatty and Lucero-Wagoner, 2000; van der Wel and van Steenbergen, 2018). It is also interesting to note that training typically leads to decreased (not increased) brain activation - plausibly, by increased knowledge of task contingencies and the ensuing decrease of the metabolic costs associated with task-related information processing (Solopchuk et al., 2017; Wiestler and Diedrichsen, 2013).

### 3.3 The costs of counteracting priors or default policies

Following the same reasoning as above, our framework predicts also that tasks that require *counteracting deep priors* or *default policies* would be particularly demanding (see Figure 5). Indeed, if priors are not just uninformative, as assumed in the previous paragraph, but also counter-productive, assuming strong statistical relationships that are no longer true, the KL divergence with the true joint distribution will be even larger.

**Figure 5.**
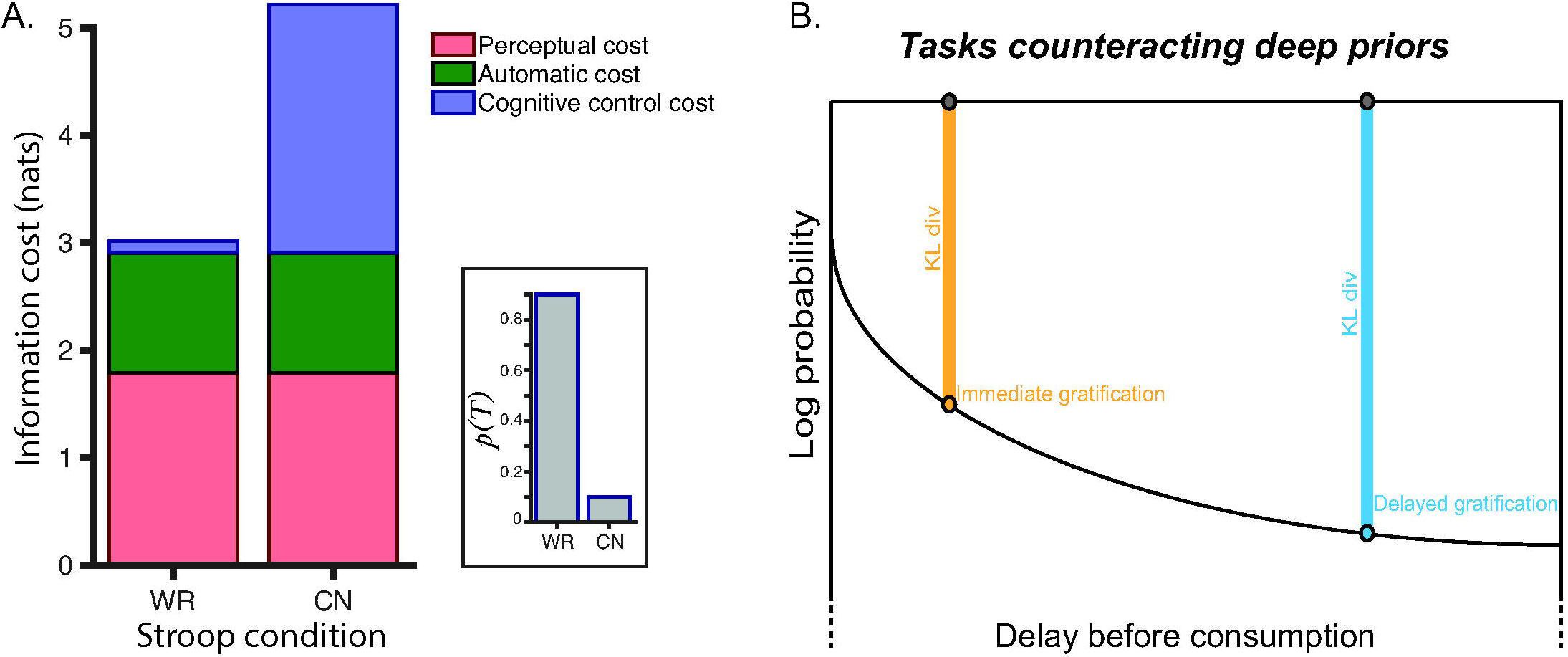
A. Simulation of Stroop task. The different sources of information cost are shown for word-reading (WR) and colour-naming (CN) Stroop trials. Units of information cost are in nats, since they were computed with natural logs. The inset shows the assumed probability of the two task contexts. The larger probability of word-reading task leads to biased automatic output *p(y*|*z)*, which explains the difference in the information cost between the two conditions. The present simulation assumes three possible colours. The different values are computed as follows (here z is assumed to be equal to x, representing the different stimulus configurations): *p*_0_ (*T*) = {.9,.1}, while *p*_opt_ (*T*) = {1,0}for the word-reading task and {0, 1} for the colour reading task.

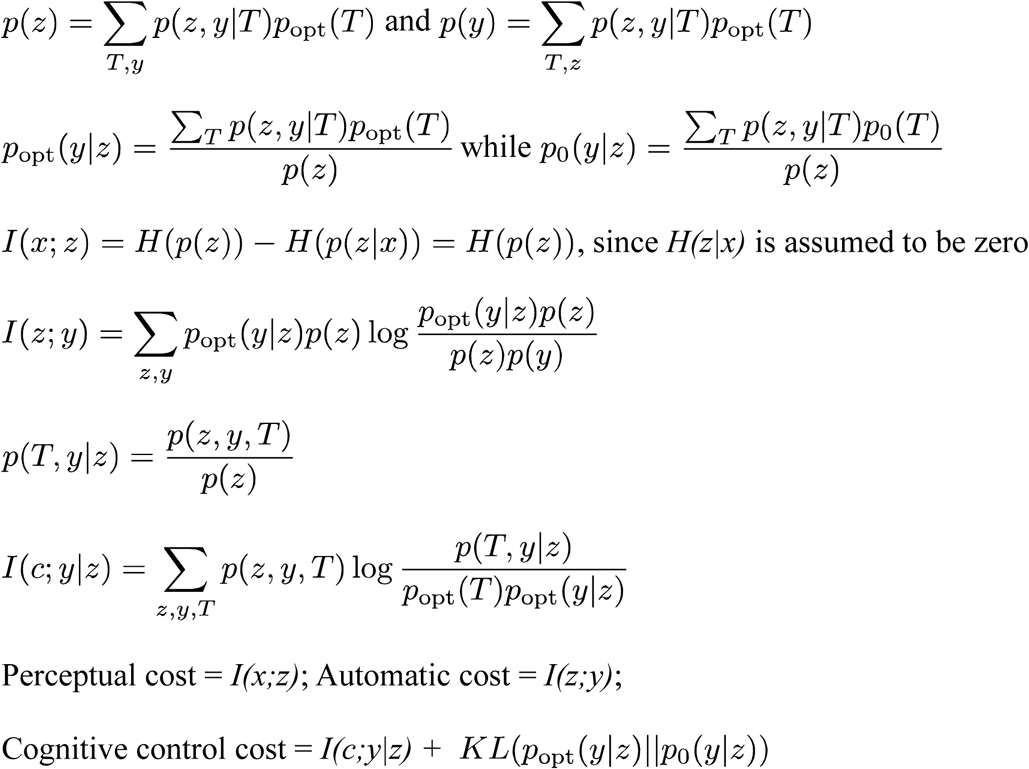 Perceptual cost = *I(x;z)*; Automatic cost = *I(z;y)*; Cognitive control cost = *I(c;y*|*z)* + *KL* (*p*_opt_ (*y*| *z*) ‖ *p*_opt_ (*y*| *z*)) B. Informal illustration of the marshmallow test, which is an example of a task requiring to counteract default policies and deep priors (Mischel, 2014). Children are told that they will receive 2 marshmallows if they don’t eat the one in front of them. The prior (and/or default policy) associated to consumption of high carbohydrate food is skewed in favour of immediate consumption, explaining, informally, why restraining from eating the marshmallow requires self-control and effort.

Similarly to the case of unfamiliar tasks exposed in section 3.2, the extra cost of using the wrong priors can be formalized as:

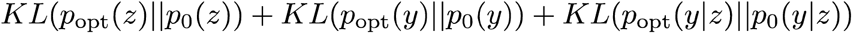

However, whereas the priors *p*_*0*_ are easy to estimate in the case of novel tasks, since they are simply non-informative (e.g. uniform distributions), in the present case, they can take many different forms and will have usually to be inferred on the basis of participants’ behaviour.

In the example of the Stroop task, the prior on the response issued from the automatic process *p(y*|*z)* will be counterproductive in cases of incongruent colour naming task. One plausible way to model this incorrect prior is by taking the marginal 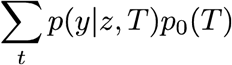 in which the probability of the word-reading context *p*_*0*_*(T=word-reading)* largely dominates the probability of the colour-naming context. This is incorrect in the sense that in the context of the Stroop task, word-reading is no longer more likely than colour-naming. This results in a marginal which favours word-reading responses, leading to extra processing costs in incongruent colour-reading trials (see Figure 5 A & B). The advantage of this approach is that cost depends only on the task structure *p(x,y)* and on the assumed prior context probability *p*_*0*_*(T)*. Here, for simplicity, we considered point estimates for *p*_*0*_*(T)* but Dirichlet distributions could be considered, with the advantage of associating a precision to the belief on context probabilities. Very high precisions would be associated with very rigid beliefs on context probabilities, leading to costs that would be relatively insensitive to training.

The present framework can explain why counteracting habits – or default responses to environmental stimuli – is so costly. Habitual behaviour is characterized by fast, automatic processing, low effort and lack of flexibility (Kahneman, 2011; Moors and De Houwer, 2006; Schneider and Chein, 2003). Under the present framework, with overtraining, the encoding of task-specific information follows so closely the statistical task structure that all the task-irrelevant information gets ignored. The ensuing cognitive processing is thus extremely efficient but also crucially dependent on the particular task contingencies that have been learned. Expected stimuli and their associated actions have very large prior probabilities *p*_*0*_*(z), p*_*0*_*(y), p*_*0*_*(y*|*z)* and are therefore encoded with minimal cost, while unexpected stimuli or actions have very low prior probabilities and are therefore very costly to encode. This implies that a person following habitual policies has lower costs to engage in familiar tasks but higher costs to engage in novel tasks. This impact of familiarity on cognitive cost could explain why novel environments (e.g. new places, new languages, new people, etc.) are generally described as being more fatiguing than familiar ones, while natural, familiar environments would have, on the contrary, restoring effects (Kaplan and Berman, 2010).

Priors can also have a deeper meaning from the perspective of the active inference framework (Friston, 2010). Under that view, agents are equipped with hierarchical generative models and perform Bayesian inference; and have the general objective to minimize their free energy or, with some simplifications, their surprise - or the discrepancy between what they expect, based on their beliefs, and what they sense. Importantly, they can minimize their surprise in two ways: by changing their beliefs to make them more similar to what they sense about the world (i.e. perceptual processing) or by changing the world to make it more similar to their prior beliefs (i.e. using actions to fulfil one’s own expectations). This duality is possible if one considers that active inference agents are hierarchically organized. While hierarchically lower prior beliefs might faithfully adapt to the external world, hierarchically deeper priors would prescribe what states an agent should achieve by acting (Friston et al., 2012; Pezzulo et al., 2015). These latter, deeper priors hence play the role of goals and motivational factors that are relatively less permeable to learning (i.e., in Bayesian terms, they have very high precision or inverse uncertainty) because they are key to survival. Indeed, a key statement of active inference is that biological agents need to minimize their long-term surprise in order to survive; if one thinks of these deep priors as describing the “good” states in an agent’s ecological niche, minimizing surprise means that the agent should attempt to remain always close to these states. One example of deep (and perhaps hard-coded) prior is a homeostatic drive, such as the prior probability of body nutrients being within acceptable physiological range. A discrepancy (prediction error) between such deep prior (e.g., be satiated) and the current interoceptive sensations (e.g., feeling hungry) would not lead to the revision of the prior - since the prior is largely impermeable in virtue of having high precision. Instead, the prediction error would steer a cascade of predictions about the conditions that might restore body nutrients (e.g., consuming food), which in turn would steer an adaptive policy or action sequence to fulfil these predictions (e.g., open the fridge and take some food). This formulation makes it apparent that a hungry active inference agent would assign a high probability to (predicted) states and policies associated to consuming food. Since the information theoretic perspective on the costs of cognition advanced here assumes that counteracting such high-probability states (or equivalently, pursuing low-probability states) has high information costs, any task that necessitates counteracting deep priors or their ensuing policies should lead to large information and cognitive costs (see Figure 5 C). This perspective may help to explain the effortful nature of self-control (Kool et al., 2010), which consists precisely in going against natural biases, as investigated in the large, but still very controversial literature on ego depletion (Hagger et al., 2016, 2010; Job et al., 2010; Kurzban et al., 2013; Muraven and Baumeister, 2000). The same arguments, based on strong (deep / homeostatic or shallow / habitual) priors can help understand the phenomenology associated to some pathological situations, such as Tourette syndrome and obsessive-compulsive disorders. These and other syndromes have been associated to strong priors that are, however, maladaptive and lead to inappropriate behaviour (Adams et al., 2013; Friston et al., 2014). Patients suffering from these disorders typically describe being able to overcome their (habitual) tics (Delorme et al., 2016) or compulsive behaviour but at the price of tremendous cognitive effort (Kawohl et al., 2009).

### 3.4 The costs of task switching and dual tasks

Another classic cause of cognitive effort is *task switching* (see Figure 6). Changing from a task set to another is associated to a significant cost in performance, usually measured as increased reaction time (Wylie and Allport, 2000). Task switching is also accompanied with a subjective cognitive effort cost (Apps et al., 2015; Kool et al., 2010) and leads to cognitive fatigue (Borragán et al., 2017). Reaction time costs in task switching have already been modelled by means of information theoretic approaches (Cooper et al., 2015). Here we propose to frame switching costs within our general framework. This can be done by considering that tasks A and B, for example, correspond to two different contexts *T* associated with different probabilities *p(T)*. Following one trial in task A, *p(T=A)* increases and the marginal 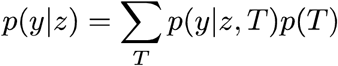 becomes closer to the correct response probabilities for task A: *p*(*y*|*z, T* = *A*) However, and for the same reason, when the task switches to B, the context-dependent cost *I(T;y*|*z)* becomes larger. Therefore, the faster the participant learns the task structure, and the longer she/he is trained on that task, the more expensive the switch cost will be. Practically, this situation can be parametrized only by the speed of learning of the context probability *p(T)*.

**Figure 6.**
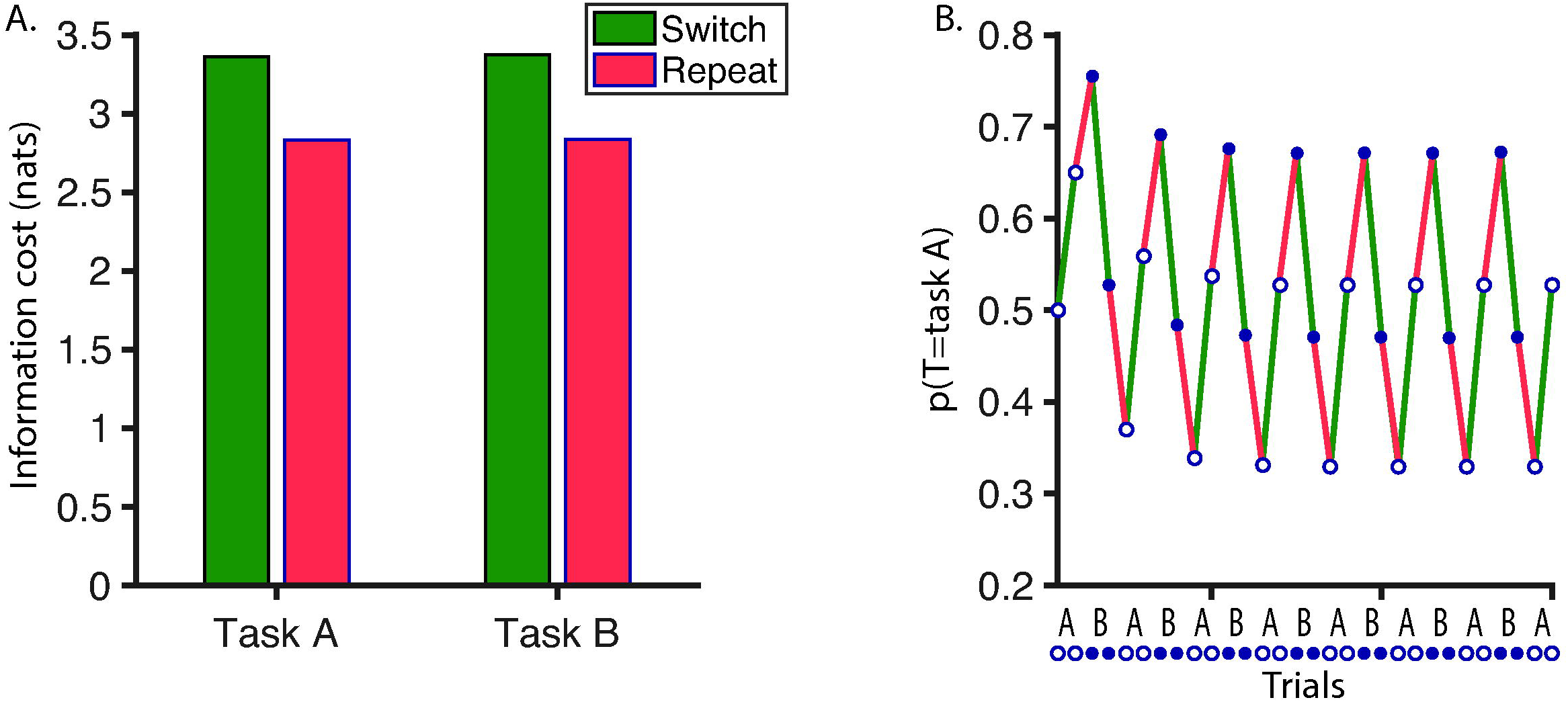
Simulation of switching task. Two tasks, A and B, alternate every other trials. Both tasks involve the same stimuli but different stimulus-response associations. A. Total information cost is shown for switch (green) and repeat trials (red). The difference between these two types of trials is entirely explained by the change in *p(T)*, shown in panel B. B. The change in the modelled *p(T=task A)* is shown as a function of trials. The experiment begins with uniform task (or context) probability distribution *p(T)*. In each trial *p(T)* is updated with a learning rate α of 0.3:

*p*_*t* +1_ (current task) = *p*_*t*_ (current task), + *α* (1 − *p*_*t*_ (current task) and *p*_*t* +1_ (other task) = *p*_*t*_ (other task) + *α* (0 − *p*_*t*_ (other task) Changes of *p(T)* following switch and repeat trials are shown in green and red, respectively.

The same arguments may apply to more mundane situations in which one is required to switch continuously between multiple tasks (multi-tasking); or to dual-task situations, in which one has either to maintain a sophisticated internal model where the probability distribution spans the contingencies of both tasks, or to rapidly and repeatedly switch between the tasks to be executed concurrently.

Interestingly, our model makes the specific prediction that switching tasks involving common stimuli (e.g. switching between colour-naming and word-reading Stroop task (Wylie and Allport, 2000)), should lead to larger costs than when switching between tasks with different stimuli, in agreement with the literature (Rubin and Meiran, 2005). This is because when different stimuli are involved, the automatic process will lead to different response probabilities for each stimulus *p(y*|*z*_*A*_) and *p(y*|*z*_*B*_), decreasing interference between the tasks. Moreover, our model also explains naturally task set inertia: the observation that interference from task A persists long after switching to task B (Allport et al., 1994). Indeed, following the switch, *p(T)* will update in each trial, getting larger and larger for task B, and smaller and smaller for task A, thereby improving progressively performance of task B.

### 3.5 The costs of controlling the rate of information processing

Another task feature that is well known to affect cognitive cost is *signal to noise ratio* (see Figure 7). When the ratio between signal and noise in sensory data is low, performance decreases, pupil size increases and subjective cognitive effort increases (Manohar et al., 2015; Sarampalis et al., 2009; Zekveld et al., 2014). Since performance and information costs are subjected to a trade-off, increasing task performance implies larger information costs. Likewise, when sensory information is immersed in high noise, and one wants to maintain reasonable performance, one needs to raise the encoding precision by decreasing the level of compression of *x* into *z* (the encoding of sensory data) and hence increase information costs by raising *I(x,z)*. Rate distortion theory provides a framework for addressing this type of problem, by describing the relation between minimal information rate (i.e. number of bits used per symbol encoded) and performance (or distortion) in a given information processing system (Shannon, 1959; Sims, 2016). Simply put, in order to minimize information cost, one should compress input data in order to discard information that is irrelevant to the task. For a given task, null distortion (i.e. perfect performance) is associated with a minimal information rate, which corresponds to the entropy of the task (i.e. its average surprisal). Beyond this level of compression, relevant information is necessarily discarded and distortion starts to increase, leading to lossy compression. This trade-off is usually implemented as a parameter that constraints the capacity (or maximal mutual information) of the system (Alemi et al., 2016; Denève et al., 2017; Genewein et al., 2015; Kingma and Welling, 2013; Ortega and Braun, 2013; Tishby et al., 2000): 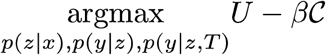, where *U* represents some utility measure and *β* is a (Lagrangian) factor that adjusts the trade-off between costs and performance. In order to apply this framework to behavioural data, this *β* parameter then has to be fit to observed performance data (Sims, 2016).

**Figure 7.**
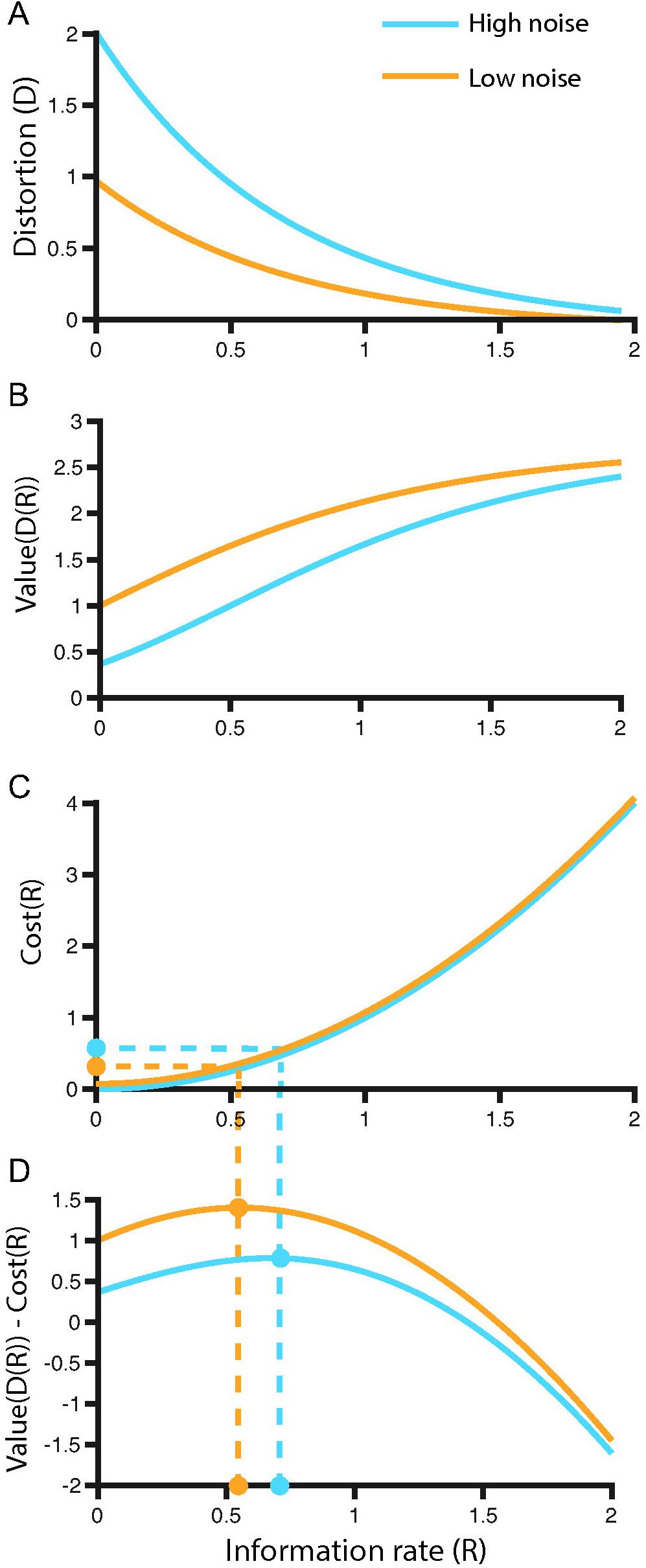
Adjustment of information rate as a function of noise levels. A. Schematic relationship between information rate and distortion in a sensory discrimination task in two conditions of noise levels. Distortion is a measure of error rate (i.e. reciprocal of performance). Assuming distortion is quantified as the mean-squared error and the signal follows Gaussian distribution with variance 2: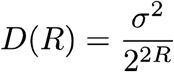. B. Utility *U* associated to distortion as a function of information rate in the 2 noise level conditions (arbitrary concave function:). *U* (*D*(*R*)) = ^*e 1*− *D*(*R*)^) C. Cost function of information rate. This cost function is independent of the noise condition, leading to overlapping curves. An arbitrary convex function was chosen:*C* (*R*) = *R*^2^. The dashed lines indicate the cost corresponding to the optimal information rates in both conditions. D. Net value, corresponding to the value associated to distortion levels, to which the cost of information rate is subtracted. The optimal trade-off between distortion and information rate should be the one associated with maximal net value, as shown with the dashed line on the plot.

This adjustability of information rate evokes the concept of task engagement and the fact that motivational factors and reward incentives can influence task performance by putting high information rate at a premium (Camerer et al., 1999). This mechanism implies some cost-benefit computation to select the optimal trade-off between the cost of information rate and the value associated with performance (see Figure 7A-D), akin to many earlier models of effort-based decision making (Chong et al., 2017; Christie and Schrater, 2015; Rigoux and Guigon, 2012; Shenhav et al., 2013; Verguts et al., 2015).

### 3.6 Summary so far

To summarize, we have offered a unitary perspective that may explain many experimental findings on cognitive effort and fatigue - by appealing to the fact that specific kinds of tasks that are known to be cognitively costly, such as novel tasks or those that require counteracting habitual policies or switching contingencies, all have high information costs associated to encoding or revising probability distributions within the generative models that support task performance. Equipped with this formalization of cognitive costs, we can now turn to their relationship to effort and their potential implementation in the brain.

## 4. Relevance for subjective effort and task avoidance

As mentioned in the introduction, the cost of cognition manifests itself behaviourally as a subjective percept of effort and as a tendency to avoid demanding tasks. Previous characterizations of the origin of cognitive effort can be classified into two broad categories. First, effort can be framed as a consequence of resource limitations such as depletable metabolic precursors (reviewed in Shenhav et al., 2017). Second, cognitive effort can be described as the phenomenological manifestation of the opportunity cost of engaging limited cognitive resources in demanding cognitive tasks (Kurzban et al., 2013). In the following, we will detail the expected consequences of the implementation in the brain of information costs, as described above. We will show that these consequences can be reconciled with the two aforementioned views on effort, while adding some novel quantitative predictions.

Before that, a methodological caveat is necessary. We have discussed how cognitive computational costs should be evaluated in terms of KL divergence between priors and posteriors. However, to apply this framework – and measures of information like surprisal, entropy and mutual information – to understand brain cognitive costs, it is necessary to assume that the brain uses an optimal strategy to encode its variables of interest. In such a code, each datapoint is represented with an average number of symbols which is proportional to the negative log of its probability (Shannon, 1948). Here probability is meant in the sense of predicted occurrence, given all the information we have at our disposal. For instance, the word “hatter” is not very common in English sentences in general but a sentence starting with “As mad as a” is much more likely to be continued with the word “hatter”. In this latter case, the word “hatter”, under optimal encoding strategy, should be encoded with small number of symbols (Lai, 2009). So, in order for the proposed framework to be applicable to brain processes, we need to assume that the brain indeed approaches such an optimal encoding strategy. This assumption is at the core of the efficient coding hypothesis (Collell and Fauquet, 2015; Harremoës and Tishby, 2007; Laughlin, 2001; Sims, 2016; Tkačik and Bialek, 2014; Wei and Stocker, 2015). Efficient coding was initially described as a theory of redundancy reduction, according to which biological systems decorrelate sensory signals to avoid redundancy (Attneave, 1954; Barlow, 1961; Simoncelli and Olshausen, 2001). The theory has been successively extended to include other mechanisms through which neural coding adapts to the statistical structure of its environment (Simoncelli, 2003; Smith and Lewicki, 2006; Tkačik and Bialek, 2014). One crucial aspect of efficient coding is the usage of an adaptive code, in which the cost for encoding each symbol is inversely proportional to its frequency in the environment - given the agent’s model of the environment (Collell and Fauquet, 2015; Fairhall et al., 2001).

An impressive body of experimental evidence has been accumulated in favour of this hypothesis (Tkačik and Bialek, 2014). Low-level vision and audition show data filtering properties and neural codes that are closely similar to predictions issued from efficient coding models (Borst and Theunissen, 1999; Gutnisky and Dragoi, 2008; Laughlin, 2001; Olshausen and Field, 2004; Sharpee et al., 2006; Smith and Lewicki, 2006). Predictability leads to diminished brain activation (Auksztulewicz and Friston, 2016; Bell et al., 2016; Carreiras et al., 2009; Garrido et al., 2013; Lieder et al., 2013; Mars et al., 2008; Meyniel et al., 2016; Overath et al., 2007; Wacongne et al., 2012) and pupil responses (Friedman et al., 1973), but increases reliability of encoding (Kok et al., 2012), in agreement with the idea that predictable stimuli, carrying little information, are encoded more economically. Along the same line, decreased brain activation following training (Chen and Wise, 1995; Solopchuk et al., 2017; Toni et al., 1998; Wiestler and Diedrichsen, 2013) suggests that training decreases metabolic cost by allowing learners to leverage task statistics to optimize brain representations, while increasing the quantity of information being processed. Even though these pieces of evidence do not yet allow us to consider the brain usage of optimally efficient coding as an established fact, we will assume here that the brain indeed uses an efficient code since this assumption remains a pre-condition for our framework to be applicable.

Another important question, if one is to apply the present information theoretic perspective to the brain, concerns the ways different brain areas are taxed by costs. The brain is massively parallel, and while the total information cost of a task can be evaluated, it remains to be determined how the cost affects (and is shared between) specific brain areas and individual neurons within these areas. One possible starting point would be to assume that brain networks supporting perceptual and decision processes would be taxed by the three kinds of costs considered in our proposal; namely, the costs associated to a perceptual stage, in which the sensory input *x* would be represented through an internal variable *z;* the cost of the automatic stage, in which the stimulus leads to a response independently of context; and the costs associated to context-dependent response selection, which would amount to updating *p(y*|*z)* to the final response *p(y*|*z,T)*. While this is certainly a simplification (as perceptual and decision processes are not segregated in the brain), it would provide a first rough set of hypotheses on the ways different brain areas would be affected by different kinds of costly tasks. However, a more complete view should consider the hierarchical organization of perception-action loops in the brain (Fuster, 1990), and relate different kinds of costs to hierarchical lower and higher stages of cognitive processing.

It is worth noting that hierarchical processing can be viewed either as serial or parallel. Serial processing has the major drawback that total information capacity of the system is equal to the information capacity of its weakest link (Genewein et al., 2015). In contrast, parallel processing consists in architectures in which the outputs from high-level processes provide priors, rather than inputs, to low-level processes (Genewein et al., 2015) and in which the information capacities of the individual parts sum up. Here, while we consider perceptual processing as a serial process (information is first stored in a variable *z* and then processed further to provide response *y*) automatic and context-dependent processes follow a parallel architecture, in which the outcome of the automatic process is fed as a prior into the context-dependent one. This parallel hierarchical architecture evokes predictive coding and active inference, which are built on this type of organization (Bastos et al., 2012; Clark, 2013; Friston et al., 2009; Pezzulo et al., 2018; Rao, 2010), or the work of Koechlin and Summerfield, which proposed such a parcellation of cognitive control costs in terms of depth of contextualization (Koechlin and Summerfield, 2007). Naturally, one can leverage the vast neuroimaging literature in order to estimate the decomposition of cognitive tasks into relevant sub-processes but studies using directly information theoretic approaches to study brain activations will be necessary to achieve better specificity (Kriegeskorte and Bandettini, 2007; Wu et al., 2017).

These caveats in mind, we can propose two (non exclusive) approaches to explain why information cost should lead to the phenomenological perception of effort and to task avoidance. The first approach is to consider that the information capacity of brain areas (i.e. the maximum of the mutual information between the data and its neural representation) is limited. Therefore, increasing the information demand necessarily leads to decreased capacity for other concurrent processes and when a given process utilizes the full capacity, larger information demands translate into longer reaction times. The second approach consists in considering that information costs have direct equivalence in terms of energetic demands. According to this view, the constraint on the system is energetic, rather than informational. This metabolic constraint leads in turn to two potential consequences on subjective cognitive costs in terms of global opportunity costs and metabolic alterations. These different points of view are detailed below (see also Figure 2, blue panel).

### 4.1 The information capacity perspective on cognitive costs

Neurons have limited information processing capacity (Schneidman et al., 2000), and so do brain areas (Marois and Ivanoff, 2005; Verghese and Pelli, 1992). Therefore, taxing of brain area capacity by a cognitive task will decrease the capacity left for other processes. The functional significance of this decrease depends on the affected brain regions. Here for brevity we focus on two brain networks: sensory cortices and the multiple demand system. The sensory cortices encompass large area, with topographic organization, in which processing of different input features or spatial locations leads to activations of different cortical regions (Purves et al., 2001). Declines in capacity in such topographic areas are easy to compensate. The situation may be different in the case of the multiple demand system: an ensemble of brain areas engaged in a large variety of tasks (Fedorenko et al., 2013), including interoceptive processing (Kleckner et al., 2017), i.e. the adaptation of behaviour to fullfil physiological needs, a function essential to survival. Thus, decrease of information capacity within this network may have more adverse behavioural consequences (and presumably more severe effort and fatigue phenomenology). This bridges the present framework with the opportunity cost view on mental effort (Kurzban et al., 2013), according to which effort must be understood as the cost of forfeiting potentially more valuable courses of action than the current task, due to taxing of limited cognitive resources.

So, when a task taxes part of the information capacity of a brain structure, its cost can be expressed in terms of the limitation it imposes on which other processes, dependent on the same structure, could be run in parallel (Kurzban et al., 2013). However, another important determinant of this opportunity cost is how long the task lasts, or in other words, its associated reaction time. This is especially crucial for tasks whose performance depends on information bottlenecks, i.e. on brain structures whose capacity is fully engaged in the task. The original findings of Hick and Hyman suggested that such capacity limit was reached for tasks as simple as stimulus-response associations (Hick, 1952; Hyman, 1953), since reaction times in their studies was a linear function of information cost. This suggests that in many circumstances, rather than as a limit on multi-tasking, information cost must be understood in terms of the opportunity cost of time (Niv et al., 2006; Payne and Bettman, 1996; Zénon et al., 2016), i.e. how long brain resources remain dedicated to the same cognitive activity (Barrouillet and Camos, 2012). In an attempt to formalize these ideas, we propose that, according to the capacity perspective, subjective costs are proportional to the maximum, across all subprocesses, of the ratio between local information costs and local capacity. Thus, subjective cost ℱof a cognitive activity could be approximated as 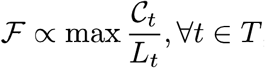, considering that the cognitive activity is composed of an ensemble *T* of subprocesses *t*, that 𝒞_*t*_ is the information cost of a specific subprocess and that *L*_*t*_ is the information capacity of the corresponding brain structure (we use *L* rather than the standard symbol *C* to avoid confusion with the cost symbol).

### 4.2 The metabolic perspective on cognitive costs

Information processing requires energy expenditure (Landauer, 1996; Ortega and Braun, 2013; Sengupta et al., 2013; Still et al., 2012). The relation between informational and energetic costs was central in early theories of efficient coding (Atick, 1992; Attneave, 1954; Barlow, 1961; Borst and Theunissen, 1999; Niven and Laughlin, 2008), which assumed that neural responses, carrying large energetic costs, impose a constraint on brain information processing capacity. Under this framework, the brain attempts to maximize the amount of (mutual) information it processes, given this fixed energetic constraint. More recent approaches have relaxed this principle by considering the constraint to be adjustable (Denève et al., 2017; Ortega and Braun, 2013; Park and Pillow, 2017; Sengupta et al., 2013), opening the door to cost-benefit adjustments of the kind exposed above (see Figure 7), which allow metabolic costs to be adjusted as a function of demands in performance (Genewein et al., 2015; Park and Pillow, 2017; Sims, 2016). So, the metabolic perspective on cognitive costs assumes that subjective effort and task avoidance have a fundamental energetic origin, and that energetic costs *E* are a function of information costs as defined above: *E*. = *f* (𝒞) Even though determining function *f* precisely is difficult, approximating it is possible with standard neuroscience techniques. For instance, tackling this question with neuroimaging in the sensory domain could be done by relating the amplitude of the haemodynamic response of brain areas (approximating energy demands) to the mutual information between their activations and their inputs.

If multiple subprocesses *t* are running in parallel the energetic cost depends on the sum over the information costs of these tasks. However, we now must add another constraint to this formula, which is that the total energetic cost across the ensemble of brain processes *B* should be constant: 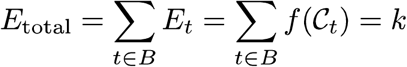

This constraint comes from the observation that global cerebral energy consumption does not vary between resting and active conditions (Lennie, 2003; Sokoloff, 2009; Sokoloff et al., 1955). Blood delivers glucose and oxygen to brain in excess of demand, such that in physiological conditions (i.e. in absence of hypoxia or hypoglycaemia), the availability of energetic precursors is not a limiting factor to cognitive activity (Brown and Ransom, 2014). However, total blood delivery to the brain is a constant that cannot be upregulated in response to cognitive demand (Brown and Ransom, 2014). Therefore, the cost of cognitive activity can hardly be explained by the need to curb total energetic consumption. It is noteworthy, however, that despite this lack of change in global energetic demand of the brain, global glucose intake increases in the brain during cognitive activity (Volkow et al., 2008). This utilization of glucose in excess of oxygen consumption is referred to as aerobic glycolysis (Vaishnavi et al., 2010). It is modulated by arousal (Dienel and Cruz, 2016) and while the function of aerobic glycolysis remains debated, it may be linked to cortical plasticity (Goyal et al., 2014) and the replenishment of glutamate and GABA reserves (Hertz and Chen, 2017). This increased glucose demand during active behaviour may appear to justify resource depletion theories of cognitive effort, according to which glucose is the main resource that puts a constraint on cognitive activity (Gailliot and Baumeister, 2007). However, this theory, and the experimental evidence on which it is based, have been put under increased criticism recently (Hagger et al., 2016, 2010; Kurzban et al., 2013; Molden et al., 2012; Shenhav et al., 2017).

Here we propose two alternative views. First, if total brain energetic consumption is constant and each brain region consumes energy in proportion to the amount of information it processes, then we must assume that neural activity tied to the execution of a cognitive task, irrespective of where it takes place in the brain, should decrease the total capacity available to other processes. This generalizes the point made in the previous section, which considered capacity as a local feature, imposed by limited information capacity of neurons and brain structures. In other words, the brain as a whole has limited capacity, due to its constant energy consumption, and allocating metabolic resources to one process leads to opportunity costs related to the reduced capacity available to other processes: 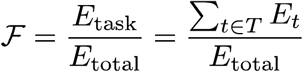, in which *T* represents all subprocesses involved in a specific task.

Therefore, increased metabolic costs in specific brain regions would entail decreased demands in other regions, which is in line with the findings that brain activations during cognitive tasks are accompanied by commensurate deactivations in resting-state brain areas (Fox and Raichle, 2007).

A second metabolic point of view on the question of why demanding tasks are avoided is that local information costs lead to progressive local metabolic alterations that accumulate over time. The rate of accumulation of these alterations 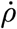 would then be a function of neural activity, itself proportional to information costs: 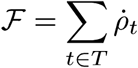

Cognitive costs would then be proxies that the brain uses to prevent these local metabolic alterations to occur. This point of view has the advantage of providing a direct link between *effort*, which would become the subjective experience of the rate of accumulation of local metabolic alterations (Tucker and Noakes, 2009), and *fatigue*, which would be the anticipation (Benoit et al., 2017) or the direct consequence of these alterations (Gergelyfi et al., 2015; Hockey, 1997; van der Linden et al., 2003).

Evidence that prolonged local brain activation leads to functional alterations supports this hypothesis (Mednick et al., 2002), but the exact nature of these alterations can only be speculated at this point. Some have proposed that glycogen reserves could deplete (Christie and Schrater, 2015), or amyloid peptides accumulate over time (Holroyd, 2015). Others have suggested that the accumulation of deviations from metabolic steady-state could lead cortical regions to enter a local sleep mode, characterized by slow-wave synchronization and associated with disturbed processing capacity (Siclari and Tononi, 2017). Confirming the existence and deciphering the nature of these metabolic alterations is an important challenge for future research on effort and fatigue.

### 4.3 The possible roles of arousal within this framework

Arousal could be defined as a global brain state characterized by the amplitude of synchronized low-frequency oscillations and sensory responsiveness (McGinley et al., 2015) and is controlled by brainstem nuclei and neuromodulators such as noradrenaline and acetylcholine (Reimer et al., 2016). An impressive amount of evidence has shown that arousal correlates with cognitive workload (Beatty and Lucero-Wagoner, 2000; Richter et al., 2016; van der Wel and van Steenbergen, 2018). Interestingly, arousal responds also strongly to prediction errors, or surprise (Ferreira-Santos, 2016; Friedman et al., 1973; Kloosterman et al., 2015; Lavín et al., 2014; O’Reilly et al., 2013; Preuschoff et al., 2011). Taken together, these two sets of observations put together are in line with the view that the modulation of arousal is ultimately a function of uncertainty (Yu and Dayan, 2003), or equivalently, of information load (entropy is uncertainty that needs to be resolved) – which permits to link nicely arousal to our proposed information-theoretic framework.

The functional role of increased arousal in response to information costs remains unclear. One possibility is that arousal mobilizes the metabolic apparatus orchestrated by astrocytes in response to neural activity (O’Donnell et al., 2012; Paukert et al., 2014), including glycogen reserves (Hertz and Zielke, 2004; O’Donnell et al., 2012), while also restricting the circulation of cerebrospinal fluid, thus limiting the capacity of the brain to eliminate potentially harmful metabolites (Xie et al., 2013). Interestingly, although arousal is a global phenomenon, its effect in cortex is believed to be restricted to active regions (Mather et al., 2016). Increased arousal would thus lead to increased metabolic rate in these active regions, while dampening activation in already less active background structures (Mather et al., 2016).

## 6. Conclusions and related work

The phenomenology of cognitive cost and cognitive effort - in terms of subjective feeling of exhaustion experienced when performing a cognitive task and its associated task-avoidance - is relatively well known. Yet, several aspects of the problem of the costs of cognition are currently debated, including the specification of what is costly in cognitive processing, how to quantify these costs and what is their adaptive value.

We have defended a view that starts from the idea that, if the brain encodes information in accordance with the principles of efficient coding, then the cognitive costs associated to task execution should be a function of the amount of information required to update priors to posterior beliefs. We have used this framework to explain the cognitive costs associated to different classes of tasks known to be subjectively demanding, and have showed that this informational perspective can provide a unitary perspective on several experimental findings in the literature. Furthermore, we have discussed how information costs could translate into cognitive effort (i.e., the subjective feeling associated to performing costly tasks). We have described three hypotheses regarding this link. First, subjective effort may arise from the usage of limited, local information capacity, chiefly in the multiple demand system, leading to opportunity costs. Second, subjective effort may be the consequence of the global limit on information capacity, caused by the constant energy consumption of the brain, also leading to opportunity costs, of a slightly different nature. Third, subjective effort could be caused by the accumulation of local metabolic alterations, whose rate is a function of information demand. In that case, effort and cognitive fatigue that ensue from long-term engagement in costly tasks could be considered as adaptive mechanisms that prevent individuals from performing activities that may have adverse consequences in the long run, i.e., activities that imply huge local metabolic demands. Importantly, these three proposals are experimentally testable and could help guiding future research in this domain.

Our perspective is coherent with several recent theories, which proposed that task avoidance stems from the (optimal) choice between potential policies while accounting for their respective cost. In this framework, cognitive effort is assumed to depend on the degree to which the task depends on cognitive control (Shenhav et al., 2017, 2013). The present work is in continuity with these earlier proposals as it relies on optimality principles and describes cognitive effort as a cost, which discounts the expected utility of a given course of actions (Apps et al., 2015; Chong et al., 2017; Manohar et al., 2015; Westbrook and Braver, 2015). However, our proposal departs from these theories as it addresses more directly the question of the *causes* of cognitive cost - and casts them in terms of information principles. In particular, the present framework attributes costs to general informational aspects of cognitive processing, rather than specifically to cognitive control. What determines cognitive cost is the amount of information to be processed and tasks involving cognitive control may be more costly in general because they typically require large information loads (Fan, 2014) on the multiple demand system (Koechlin and Summerfield, 2007; Wu et al., 2017), where the opportunity cost associated to utilization of limited information capacity is larger.

Koechlin and Summerfield have proposed an information theoretical approach to cognitive control that shares several aspects with our own framework. In their study, the total cost of selecting an action is framed as the sum of two terms (Koechlin and Summerfield, 2007). The first is an automatic, goal-independent association between stimuli and actions whose cost is measured as the mutual information between stimuli and actions: *I(x;y)*. The remaining cost of action selection *H(y)-I(x;y)* then corresponds to cognitive control and could be decomposed further into several hierarchical processes with different levels of contextualization. Similarly to Koechlin and Summerfield’s proposal (and others (Genewein et al., 2015; Pezzulo et al., 2013)), our framework assumes a hierarchical architecture in which the cognitive control process takes the outcome of the automatic process as a prior, making their contributions additive (Genewein et al., 2015). The present paper differs from this earlier treatment foremost by its focus on effort costs. In addition, we attempted at being more general, by adding the cost of perceptual processing, which is necessary to account for known behavioural results (Fan et al., 2008; Wifall et al., 2016) and by introducing rate-distortion theory. We also tried to clarify the formalism of the model by spelling out the different potential sources of cost. Finally, we also detailed the application of the framework to different types of demanding tasks.

Another proposal which relates to ours, addresses cognitive effort from a normative perspective, in which apparently maladaptive states (cognitive fatigue) constitute the adaptive response of an optimal controller that has (or feels having) low self-efficacy and limited control over one’s own environment, similar to learned helplessness in the animal learning literature (Stephan et al., 2016). Although we have not addressed the long-term consequences of being exposed to complex cognitive tasks, our model would be coherent with this proposal in assuming that prolonged expectations of poor outcomes, or poor control, would crystallize task-avoidance behaviour. In other words, an agent whose local metabolic resources are frequently depleted, could develop an adaptive task aversion, manifested, for example, as chronic fatigue syndrome.

The present framework has also some limitations that will need to be covered in future works. For example, it does not fully explain important features of switching costs, such as the fact that moving from a preferred to a less preferred stimulus-response association is less costly than the other way around (Wylie and Allport, 2000), or the fact that preparation periods, no matter how long, cannot suppress the switching costs (Wylie and Allport, 2000). It is also challenging to explain why switching between tasks that are based on similar stimulus features or similar response modalities is associated with smaller costs than switching between dissimilar tasks (Arrington et al., 2003). Explaining this finding would require the model to account for some hierarchical categorization of perceptual and response representations *z* and *y* (Zhao et al., 2017), that go beyond the scope of the present paper.

Another limitation of the current framework is that it does not fully account for adaptive (positive) aspects of costly cognitive processing; for example, engaging in (costly) information-foraging behaviour during learning. Challenging cognitive activity is not always experienced as aversive but may be even sought for its own sake (Cacioppo et al., 1984; Inzlicht et al., 2018); and on the contrary, idling or engaging in repetitive, monotonous tasks can be unpleasant (Nakamura and Csikszentmihalyi, 2002). Costly information-seeking is in apparent contradiction with our framework, but it can be explained by the mediating influence of intrinsic motivation and epistemic value (Friston et al., 2015, 2017). Complex tasks need mastery, and intrinsic motivation appears to depend in large part on the feeling of competence and autonomy, i.e. the feeling of being capable of performing a task, despite its difficulty (Ryan and Deci, 2000) - and in the active inference setting, on the necessity to improve one’s internal models (Friston et al., 2017). In other words, improving one’s internal model to minimize surprise in the future can compensate the short-term costs of investing effort in the present moment. Spontaneous engagement in demanding tasks would therefore depend on the cost-benefit comparison between immediate informational and/or metabolic costs and future needs, mediated by mechanisms of exploration, model learning or intrinsic motivation that are part and parcel of active inference (Friston et al., 2017) – and which complement nicely the information theoretic framework advanced here.

Another limitation of the present work stems from its reliance on a series of assumptions: it is valid only to the extent that those assumptions are true. First and foremost, it relies on the concept of efficient coding, according to which brain’s encoding of information makes optimal use of its metabolic resource, such that energetic cost is proportional to the entropy of the encoded information. Even though this concept is based on a large body of evidence, it cannot as yet be viewed as an established fact. Second, we assume a certain cognitive architecture, in which perceptual encoding of sensory information occurs first and is then fed into two parallel cognitive processes: one automatic and one context-dependent. We believe that this architecture is the simplest one that can account for all the categories of demanding cognitive tasks reviewed in the paper. Naturally, more complex architectures could be considered. Finally, we also base our estimation of cognitive cost on the priors used by subjects for perceptual encoding and cognitive control processing. The accuracy of the predictions made by the model will, therefore, depend crucially on the choice of those priors. A possible approach to this problem is to base the choice of the priors on observed data. Indeed, since priors are the only unknown parameters in the model, they can be fitted to participants’ behaviour (Sims, 2016).

In sum, we have reviewed an information theoretical perspective on the costs of cognition that links cognitive effort to principles of efficient coding and active inference. This framework systematizes and extends previous treatments that used information theoretic principles to explain cognitive control processes (Koechlin and Summerfield, 2007; Ortega and Braun, 2013; Sengupta et al., 2013). Furthermore, we have discussed how this framework permits to harmonize theories that link cognitive costs to capacity limitations and metabolic principles, while also producing novel empirical predictions.

## Acknowledgements

This work was supported by Fonds de la Recherche Scientifique (FNRS–FDP), Fondation Médicale Reine Elisabeth (FMRE), the Fondation Louvain and IdEx Bordeaux.

## Footnotes

1 Cost of the context-dependent process: 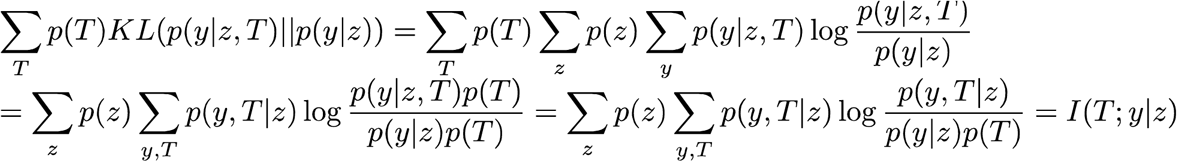

2 Derivation for the perceptual cost *I(x;z)*: 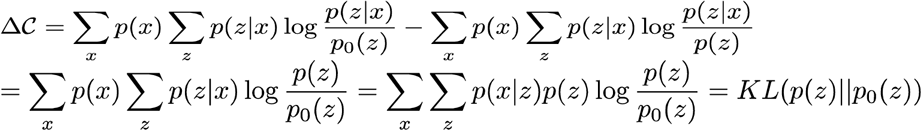

